# DNA Methylation Networks Underlying Mammalian Traits

**DOI:** 10.1101/2021.03.16.435708

**Authors:** A. Haghani, A.T. Lu, C.Z. Li, T.R. Robeck, K. Belov, C.E. Breeze, R.T. Brooke, S. Clarke, C.G. Faulkes, Z. Fei, S.H. Ferguson, C.J. Finno, V.N. Gladyshev, V. Gorbunova, R.G. Goya, A.N. Hogan, C.J. Hogg, T.A. Hore, H. Kiaris, P. Kordowitzki, G. Banks, W.R. Koski, K. Mozhui, A. Naderi, E.A. Ostrander, K.M. Parsons, J. Plassais, J. Robbins, K.E. Sears, A. Seluanov, K.J. Steinman, B. Szladovits, M.J. Thompson, D. Villar, N. Wang, G.S. Wilkinson, B.G. Young, J. Zhang, J.A. Zoller, J. Ernst, X.W. Yang, K. Raj, S. Horvath

**Affiliations:** Dept. of Human Genetics, David Geffen School of Medicine, University of California Los Angeles, Los Angeles, CA, USA; Dept. of Biostatistics, Fielding School of Public Health, University of California Los Angeles, Los Angeles, CA, USA; Zoological Operations, SeaWorld Parks and Entertainment, Orlando, Florida, USA; School of Life and Environmental Sciences, The University of Sydney, Sydney, New South Wales, Australia; Altius Institute for Biomedical Sciences, Seattle, WA, USA; Epigenetic Clock Development Foundation, Los Angeles, CA, USA; AgResearch, Invermay Agricultural Centre, Mosgiel, Otago, New Zealand; School of Biological and Chemical Sciences, Queen Mary University of London, London, UK; Fisheries and Oceans Canada, University Crescent, Winnipeg, Canada; Dept. of Population Health and Reproduction, University of California, Davis School of Veterinary Medicine, Davis, CA, USA; Division of Genetics, Dept. of Medicine, Brigham and Women’s Hospital, Harvard Medical School, Boston, MA, USA; Depts. of Biology and Medicine, University of Rochester, Rochester, NY, USA; Biochemistry Research Institute of La Plata, Histology and Pathology, School of Medicine, University of La Plata, La Plata, Argentina; Cancer Genetics and Comparative Genomics Branch, National Human Genome Research Institute, National Institutes of Health, Bethesda, MD, USA; Dept. of Anatomy, University of Otago, Dunedin, New Zealand; Dept. of Drug Discovery and Biomedical Sciences, College of Pharmacy, University of South Carolina, SC, USA; Institute of Animal Reproduction and Food Research of Polish Academy of Sciences, Olsztyn, Poland; Mammalian Genetics Unit, MRC Harwell Institute, Harwell Science and Innovation Campus, Oxfordshire, UK; LGL Limited, King City, ON, Canada; Dept. of Preventive Medicine, University of Tennessee Health Science Center, College of Medicine, Memphis, TN, USA; Conservation Biology Division, Northwest Fisheries Science Center, National Marine Fisheries Service, National Oceanic and Atmospheric Administration, Seattle, Washington, USA; Center for Coastal Studies, Provincetown, MA, USA; Dept. of Ecology and Evolutionary Biology, UCLA, Los Angeles, CA, USA; Species Preservation Laboratory, SeaWorld San Diego, California, USA; Dept. of Pathobiology and Population Sciences, Royal Veterinary College, Hatfield, UK; Dept. Molecular Cell and Developmental Biology, University of California Los Angeles, Los Angeles, CA, USA; Blizard Institute, Barts and the London School of Medicine and Dentistry, Queen Mary University of London, London, UK; Center for Neurobehavioral Genetics, Jane and Terry Semel Institute for Neuroscience and Human Behavior, University of California Los Angeles, Los Angeles, CA, USA; Dept. of Biology, University of Maryland, College Park, USA; Dept. of Biological Chemistry, University of California, Los Angeles, Los Angeles, California, USA; Radiation Effects Dept., Centre for Radiation, Chemical and Environmental Hazards, Public Health England, Chilton, Didcot, UK; Dept. of Biological Sciences, University of Manitoba, Winnipeg, Canada; Peromyscus Genetic Stock Center, University of South Carolina, SC, USA; Institute for Veterinary Medicine, Nicolaus Copernicus University, Torun, Poland; Dept. of Genetics, Genomics and Informatics, University of Tennessee Health Science Center, College of Medicine, Memphis, TN, USA; Dept. of Psychiatry and Biobehavioral Sciences, David Geffen School of Medicine at UCLA, Los Angeles, CA, USA

**Keywords:** DNA methylation, mammals, evolution, lifespan, aging, phyloepigenetic tree

## Abstract

Epigenetics has hitherto been studied and understood largely at the level of individual organisms. Here, we report a multi-faceted investigation of DNA methylation across 11,117 samples from 176 different species. We performed an unbiased clustering of individual cytosines into 55 modules and identified 31 modules related to primary traits including age, species lifespan, sex, adult species weight, tissue type and phylogenetic order. Analysis of the correlation between DNA methylation and species allowed us to construct *phyloepigenetic* trees for different tissues that parallel the phylogenetic tree. In addition, while some stable cytosines reflect phylogenetic signatures, others relate to age and lifespan, and in many cases responding to anti-aging interventions in mice such as caloric restriction and ablation of growth hormone receptors. Insights uncovered by this investigation have important implications for our understanding of the role of epigenetics in mammalian evolution, aging and lifespan.

## Introduction

The oldest common ancestor of all mammals lived approximately 300 million years ago, and over time diversified into more than 6,400 species that constitute the monotreme, metatherian, and eutherian lineages ^1^. These species exhibit a great diversity in traits such as maximum lifespan, adult body weight, brain/body ratio, locomotory modes, sleep/wake cycles, diet, and social behaviors. The lineage that eventually led to the emergence of the first modern humans, *Homo erectus*, about two million years ago, with the acquisition of a larger brain, higher cognitive abilities, and several unique physical and social characteristics ^2^. Comparative epigenomics is an emerging field that aims to combine epigenetic signatures and phylogenetic relationships to improve our understanding of gene-to-trait function ^3,4,5^. Even though genomic regulatory regions are under sequence constraints ^6^, epigenome evolution in these conserved regions seems to correlate with transcription in mammals ^3^, potentially regulates fitness and facilitates selection. A recent study even suggested that DNA methylation marks in regulatory sequences partially relates to the phylogenetic differences of animals ^5^. Previous DNA methylation studies in mammals were limited with regard to the sample size (relatively few DNA samples and few species) and the measurement platform (low sequencing depth at highly conserved stretches of DNA). To address these challenges, we profiled 176 mammalian species across 63 tissue and organ types using a methylation array platform that provides over 1k fold sequencing depth at highly conserved cytosines. This unique dataset could robustly identify co-methylation networks that underlie important species characteristics and allowed us to develop tissue-specific *phyloepigenetic* trees that match the topology and branch lengths of the mammalian phylogeny.

### DNA methylation networks relate to individual and species traits

We used a custom methylation array platform (HorvathMammalMethylChip40) that profiles 36 thousand CpGs with flanking DNA sequences that are mostly conserved across mammalian species^7^. With this mammalian array, we generated a large-scale DNA methylation dataset, which consists of methylation profiles from 11,117 samples derived from 63 tissue types of 176 different mammalian species **(Table S1)**. Of these 36,000 cytosines, 14,705 mapped to most eutherians and a subset of 7,956 also mapped to marsupials. Although analysis of individual cytosines can be highly effective, it ignores co-methylation relationships and clustering patterns that are associated with species traits. To address this, we used weighted correlation network analysis (WGCNA) ^7^ to cluster CpGs with similar methylation dynamics into co-methylation modules and summarize their methylation profiles using “module eigengenes”, defined as the first principal component that positively correlates with methylation changes in each module. WGCNA also identifies intramodular hub CpGs, which are cytosines with the highest correlation within each module. As such, eigengenes and intramodular hubs are mathematical representations of a module. The respective eigengenes of these modules were subsequently used to identify their potential correlations with various traits within and across mammalian species, such as chronological age, sex, maximum lifespan, age at sexual maturity, adult body weight and other characteristics for which the corresponding information are available.

In total, nine module networks were constructed, which differed with respect to the underlying CpGs, tissues, and how they relate to different species (**Extended Data Fig.1**). The eutherian network (Net1) was formed from 14,705 conserved CpGs in 167 eutherian species (**Fig. 1a**). This network showed the strongest association with various traits and will be the main focus of the following sections. The Mammalian network (Net2) consisted of eutherians plus nine marsupial species, reducing the number of cytosines to 7,925, which are shared between eutherians and marsupials **(Fig. 1b**). This subset network was used to identify marsupial-related modules. To identify the most conserved modules across species and tissue types, we developed seven consensus networks (see methods). These networks can be interpreted as a meta-analysis of methylation networks to identify consensus modules that are preserved across mammalian tissues (**Extended Data Fig.1**).

**Fig 1.**
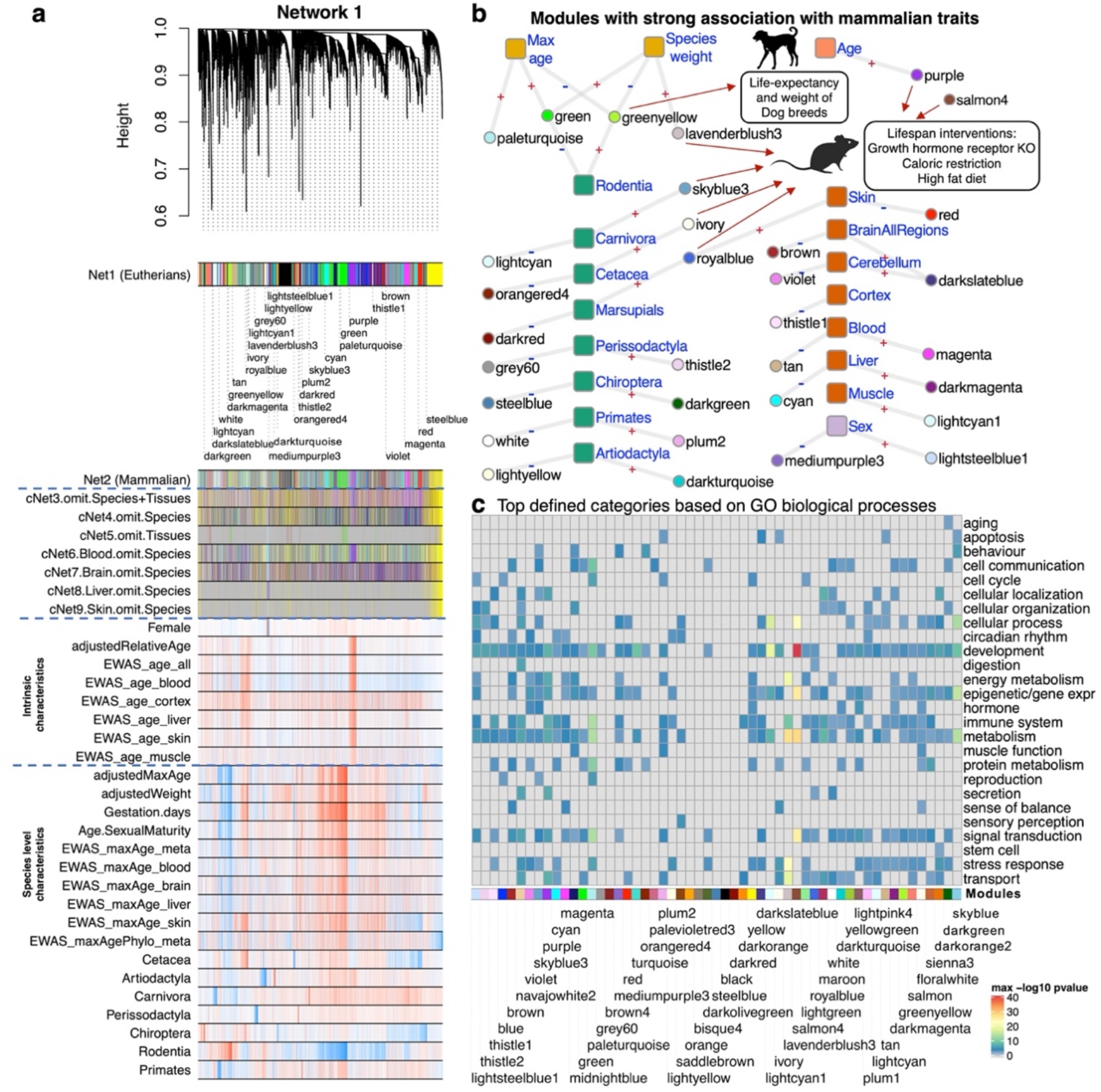
DNA methylation network relates to species and individual characteristics in mammalian species. **a**, the WGCNA network of 14,705 conserved CpGs in 167 eutherian species (Network 1). The data include around 63 tissue types, from all age ranges of most of the species. The identified modules related to species characteristics (e.g. phylogenetic order, maximum lifespan), or individual sample characteristics (e.g. tissue type, age, sex). Network 1 modules were compared to eight additional networks: based on subsets of probes that map to eutherians and marsupials (**Extended Data Fig. 1**); and seven consensus networks based on species and tissues (**Extended Data Fig. 1**). The modules with strong associations with species and sample characteristics were labeled below the dendrogram. **b**, summary of the modules that showed strong association with species and individual sample characteristics. Analyzed traits: age, sex, tissue type, species max age, species average adult weight, and different mammalian orders. The edge labels are the direction of association with each trait. **c**, Top defined functional biological processes related to network 1 modules. The gene level enrichment was done using GREAT analysis and human Hg19 background limited to 14,705 eutherian probes. The biological processes were reduced to parent ontology terms (**Extended Data Fig. 6**), and the larger ontology category was defined manually for this summary heatmap. Detailed enrichment results are reported in the supplementary excel file. Images of animals are from Phylopic (“http://phylopic.org“) or Wikimedia, which are under public domains or CC BY 3.0 license (“https://creativecommons.org/licenses/by/3.0/“).

The eutherian network (Net1) clustered the 14,705 CpGs into 55 modules **(Fig. 1a)**. These modules were derived from unsupervised clustering of cytosine methylation levels, and were labelled by colors per the convention of WGCNA (**Fig. 1a)**. The smallest module (lavenderblush3 color) consisted of 33 CpGs, while the largest (turquoise module) had 1,864 CpGs. As information on phylogenetic order, tissues, lifespan, age, sex and weight of each data point were available, it allowed us to directly assess if any of the modules were enriched for these characteristics. Of the 55 modules, 31 were found to be related to specific species or biological characteristics (linear regression p<10^−200^) (**Fig. 1b; Extended Data Fig.2; Table S3**). Specifically, 16 modules were related to phylogenetic orders. Eleven other modules related to tissue type, two modules to sex, one module to age, two modules to maximum lifespan, and two modules to average adult species weight. Some modules were simultaneously related to several characteristics. For example, the two Rodentia modules (green, and yellowgreen) were also related to species lifespan and weight. One of the marsupial modules (royalblue; identified from Net2) was also related to eutherian skin. For ease of comprehension, modules were labelled with the trait and direction of relationship by superscript +/- signs. For example, the green module was designated as LifespanWeight^(+)^Rodentia^(-)^ module The plus sign in the superscript indicates that the cytosines in the green module tend to be hypermethylated in long living and heavy species. The minus sign indicates that the green module CpGs tend to be hypomethylated in rodents. Collectively, the WGCNA results demonstrate that numerous species-specific primary traits are manifested at the level of DNA methylation; a feature that was discovered here through an unsupervised clustering of related cytosines.

Beyond association with phenotypic traits, we sought to uncover what the modules represent at the cellular and molecular levels. We identified genes that are adjacent to cytosines within the clusters and ascertained the molecular and cellular activities that are associated with them. In general, the 500 top hub CpGs of the modules were adjacent to genes associated with a wide range of biological processes including development, immune system, metabolism, reproduction, stem cell biology, stress response, aging, or several signaling pathways (**Fig. 1c**). Thus, these modules, which were derived independent of any prior biological information, are a rich source of information on the underlying biological processes that are associated with characteristics and phenotypic differences between species. Modules that relate to specific phylogenetic orders, or even species, are expected to be valuable starting points for experimental interrogation of mammalian evolution. The modules with no relationship to any analyzed traits are still valuable for future studies. For example, the yellow module with 895 CpGs is the largest consensus module that is preserved in all mammalian tissues but did not relate to any of the primary traits (**Fig. 1a**). Hub CpGs of this module are found to be adjacent to genes involved in mRNA processing, cell cycle, melanocyte development and circadian rhythm (**Table S3**). The prediction of the involvement of cytosines in the yellow module to circadian rhythm was tested in using a perturbation experiment mice. After 12 months of light pollution during night time, only the yellow module experienced a significant increase in liver methylation levels (**Extended Data Fig.2b**).

### Evolution and DNA methylation

The evolutionary divergence of three main mammalian clades (eutherians, marsupials, monotremes) occurred over approximately 250 million years (**Fig. 2a**). Many phylogenetic trees of mammals have been constructed; based initially on morphology and later on DNA or protein sequences. Prior to this study, it was unknown whether there are stable and consistent species differences in DNA methylation levels at conserved CpG sites that could allow the construction of what could be termed as a *phyloepigenetic* tree. The DNA methylation profiles at our disposal presented a unique opportunity to address this issue. To avoid confounding by different tissue types, we constructed tissue-specific *phyloepigenetic* trees. Strikingly, the resulting *phyloepigenetic* trees parallel the evolutionary distances among taxa established genetically for phylogenetic trees (**Fig. 2b; Extended Data Fig.3**). This holds true for *phyloepigenetic* trees constructed from different tissues (blood, liver and skin), as well as those from hierarchical clustering of modules based on order and species (**Fig. 2c, d)**. This result was expected as the mammalian methylation array is designed to measure methylation in highly conserved genomic sequences. The close relationship between phylogenetic and phyloepigenetic trees reflects an intertwined evolution of genome and epigenome that mediates the biological characteristics of the different mammalian species.

**Fig 2.**
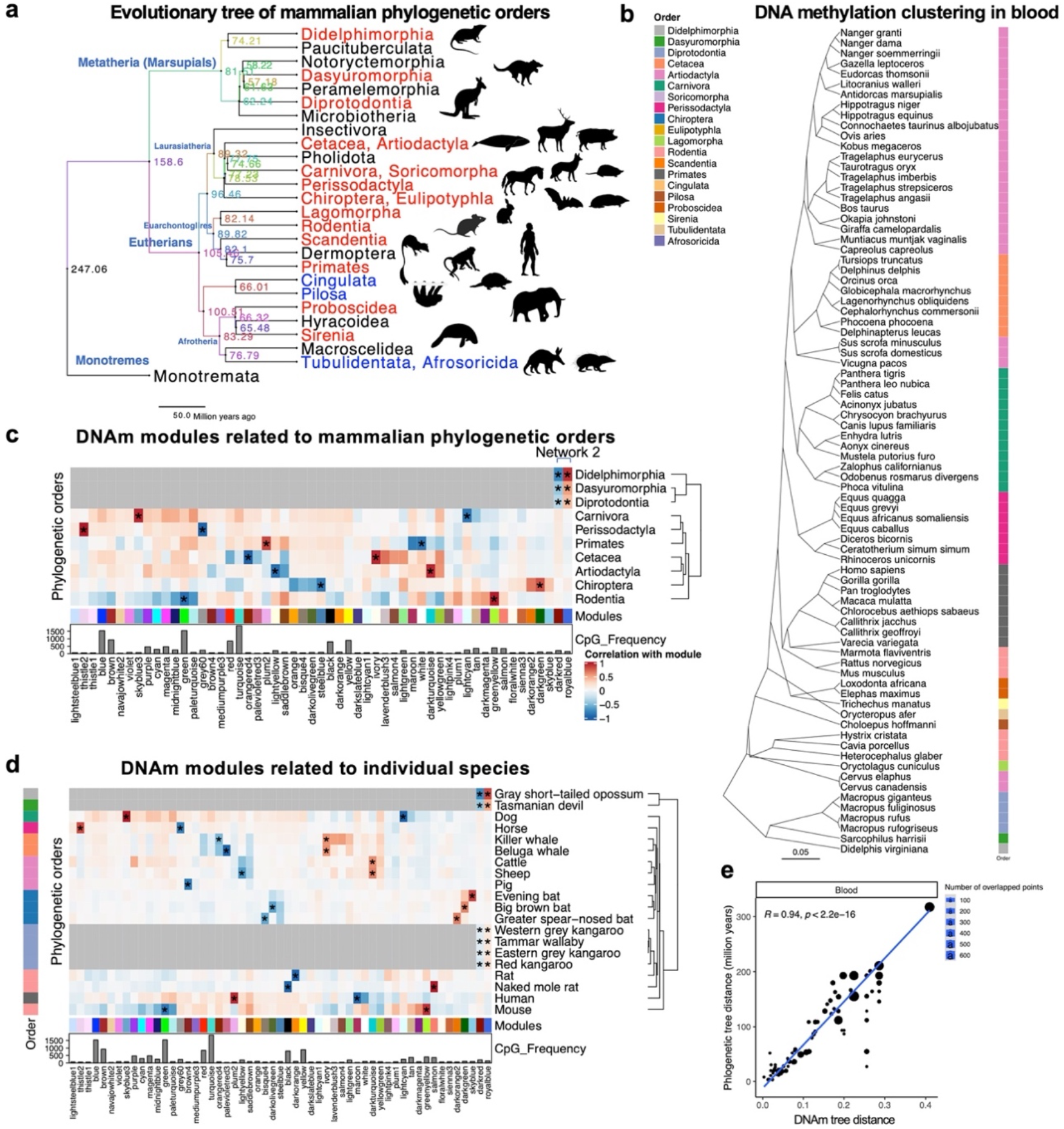
DNA methylation networks parallels the evolutionary tree in mammals. **a**, the schematic representation of the evolutionary tree in mammals ^54^. The numbers indicate the time of evolutionary divergence (million years ago) between different orders. The red and blue phylogenetic orders are included in this study. The blue indicates small sample size (<4) in our study. **b**, hierarchical clustering of DNA methylation profiles highly matches the evolutionary distance of mammalian species (**Extended Data Fig.3a,b**). Distances: 1-cor. **c**, Heatmap of phylogenetic order specific modules. * indicated the top two modules related to each phylogenetic order with minimum absolute correlation of 0.5. The marsupial modules were identified in Network 2, which was based on the conserved CpGs in both eutherians and marsupials. **d**, DNA methylation modules associated with individual species. The top two modules for each species are labeled by *. The marsupial modules are identified in network 2. The color code of the rows shows the phylogenetic order of each species. The rows are clustered based on hierarchical clustering of Euclidean distances and complete method. The species with weak module association were not shown in the heatmap. **e**, The correlation distances of hierarchical clustering of blood DNA methylation and evolutionary tree. Additional analyses are reported in the supplement (**Extended Data Fig. 4**). Images of animals are from Phylopic (“http://phylopic.org“), which are under public domains or CC BY 3.0 license (“https://creativecommons.org/licenses/by/3.0/“).

### Maximum lifespan and average adult weight

To further explore the association of modules with weight or lifespan, we categorized modules according to how they relate to maximum lifespan and/or average adult weight of the species. A single module positively associated exclusively with lifespan: Lifespan^(+)^ (paleturquoise, 87 CpGs), while another with adult weight in a positive direction: Weight^(+)^ (lavenderblush3, 33 CpGs). Interestingly, two modules relate simultaneously to max lifespan *and* weight: LifespanWeight^(+)^Rodentia^(-)^ (green, 1542 CpGs), LifespanWeight^(-)^Rodentia^(+)^ (greenyellow, 398 CpGs) (**Fig. 3a,c**). The LifespanWeight^(+)^Rodentia^(-)^ (green module) is positively correlated with maximum age (r = 0.70, p<10^−200^) and adult weight (r = 0.57, p<10^−200^), i.e. the underlying cytosines tend to be hypermethylated in long-lived and larger species. This module is enriched with genes (e.g. *GPATCH2L, NOVA2*, and *TSHZ2*) involved in development, immune system and Wnt signaling (**Table S3**). Ingenuity pathway analysis (IPA) indicates HDAC4, POU4F2, and RUNX2 as upstream regulators. The LifespanWeight^(-)^Rodentia^(+)^ (greenyellow) module correlated negatively with maximum lifespan (r = -0.71, p<10^−200^) and adult weight (r=-0.51, p<10^−200^). This negative association with lifespan remained even after adjusting for phylogenetic relationships or in marginal analysis of mean methylation per species (**Extended Data Fig.4i, Fig.3b,d**). Functional enrichment analysis of its hub genes (e.g. *TLE3, HMGB1*, and *TBCA*) indicates the involvement of translation, RAS signal transduction, adipogenesis, Wnt signaling, epithelial-mesenchymal transition, and calcium signaling. IPA analysis highlighted POU5F1, NANOG, SOX2 and POU4F2 proteins as potential upstream regulators with roles in stem cell biology. Lifespan^(+)^ (paleturquoise, max-age r = 0.63, p<10^−200^) and Weight^(+)^ (lavenderblush3, adult weight r = 0.53, p<10^−200^) are smaller modules that are related to protein and nucleotide metabolism respectively. The identified modules and other potential weight-related modules are shown in **Fig. 3d**.

**Fig 3.**
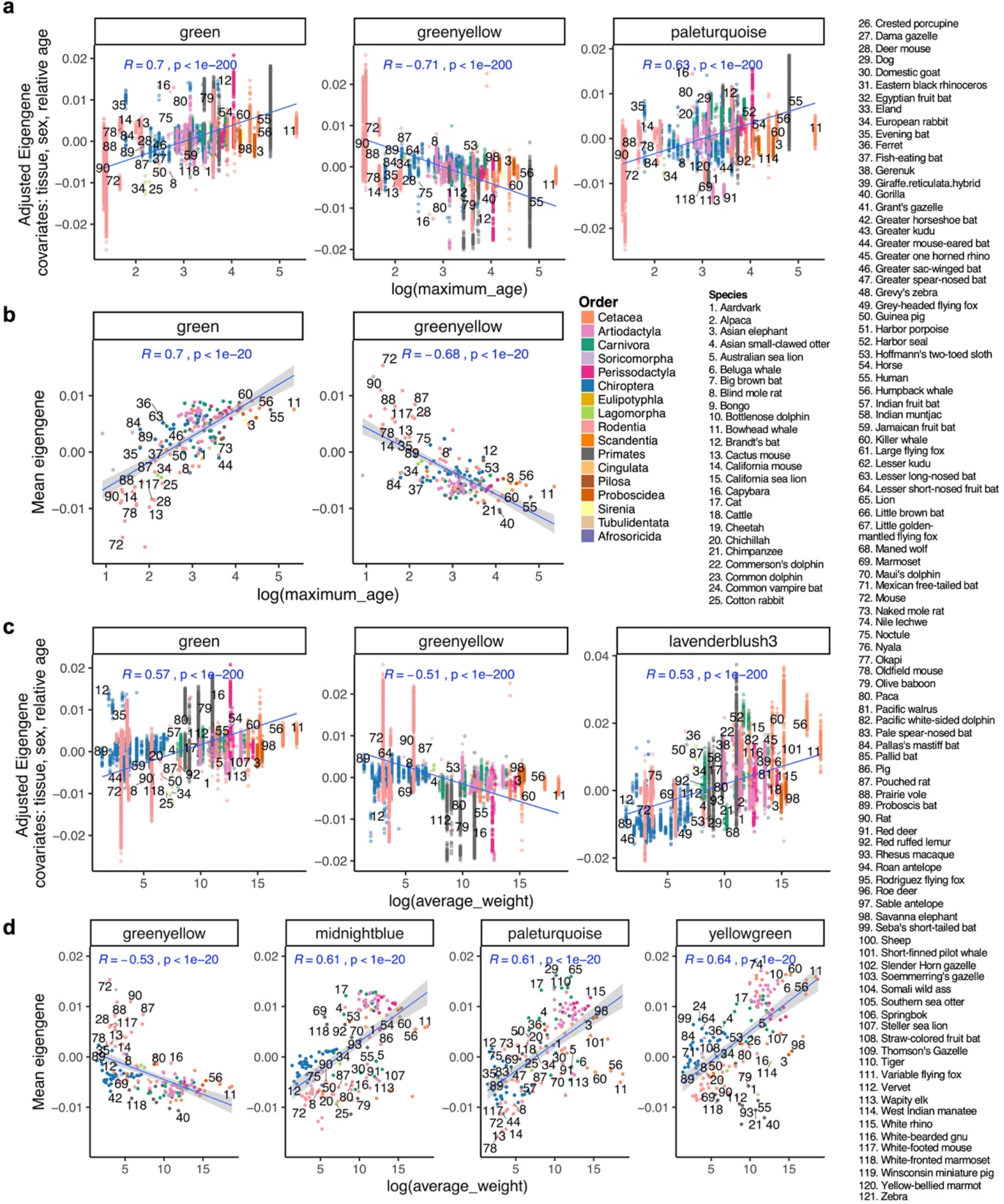
Modules with strong association with maximum age and average adult weight of the species. Associations were tested using both multivariate models and marginal association of mean eigengene. **(a)** Modules associated with log maximum age in the multivariate module (p<10^−200^). covariates: relative age, tissue, and sex. **(b)** Modules associated with log maximum age in marginal association (p<10^−20^). **(c)** Modules associated with log weight in the multivariate module (p<10^−200^). covariates: relative age, tissue, and sex. **(d)** Modules associated with log maximum age in marginal association (p<10^−12^). Eigengene: 1^st^ principal component of the module that positively correlates with DNA methylation levels.

### Consensus modules relate to tissue type, age, and sex

The mammalian methylation dataset is an atlas of 63 tissue-types, cell-types, and tissue regions from a wide age range of mammalian species. This allowed us to create a tissue module atlas and gain biological insights into DNA methylation-based mechanisms of tissue differences (**Fig.4a**). We related the module eigengenes to the following sample characteristics: tissue type, sex, and relative chronological age at the time of tissue collection. DNA methylation could distinguish the tissues with large sample sizes such as blood, skin, liver, muscle, cerebellum, cerebral cortex, and brain. The tissue modules were largely preserved in individual phylogenetic orders (**Fig.4a, Table S3, Extended Data Fig.5c**). For example, the top blood modules in different phylogenetic orders were Blood^(+)^ (magenta) and Blood^(-)^ (tan) modules. Both modules are enriched with immune system genes such as those for B cell maturation (**Table S3**). The cerebellum modules (violet, darkslateblue) on the other hand, are enriched for genes related to neuron development, projection, and differentiation pathways. Thus, tissue-specific modules corroborate the biological differences between tissue types.

Sex was strongly associated with two modules: Female^(+)^ (lightsteelblue1, hyper-methylated CpGs in female) and Female^(-)^ (mediumpurple3, hypo-methylated in female) (**Fig.4b**). Unsurprisingly, these modules consist mainly of X-chromosome CpGs (**Fig.4c**). The only autosomal CpG is located on the *POU3F2* exon in human chromosome 6. This gene showed sex-related differential methylation in primates ^8^, and is also indicated to contribute to maternal behavior differences in mammals ^9^.

We correlated the module eigengenes with two different measures of chronological age: age and *relative* age, which is defined as the ratio between age of the organism and maximum lifespan of its species (the relative age of a 61-year-old human is 0.5, as recorded maximum human age is 122.5). The purple module (denoted subsequently as RelativeAge^(+)^ module) exhibits a significant positive correlation (r = 0.37, p<10^−200^) with the relative age of all mammalian samples (**Fig. 4d**). This module also relates to the chronological age of the animal at the time of sampling (**Extended Data Fig.4f**). The RelativeAge^(+)^ module was corroborated in different consensus networks including 57 species-tissues, 35 species, and 27 species blood networks. Thus, the RelativeAge^(+)^ module is among the most conserved modules in mammalian tissues. The RelativeAge^(+)^ module contains 470 CpGs with hub genes such as *TFAP2D, CTNNA2, POU3F2, TFAP2A*, and *UNC79*. IPA implicates POU5F1, NANOG, SHH, KAT6A, and SOX2 proteins as putative upstream regulators. Functional enrichment of this module highlighted embryonic stem cell regulation, axonal fasciculation, angiogenesis, and diabetes-related pathways (**Table S3**). The CpGs in this module were adjacent to Polycomb repressor complex 2 (PRC2, EED) targets and H3K27me3 regions (**Extended Data Fig.7b**). The top hits from EWAS of age in mammals ^10^ are also enriched in PRC2 targets and similar biological processes as the RelativeAge^(+)^ module. Additionally, we carried out an overlap analysis between module genes and genes implicated by available large-scale genome-wide association studies (GWAS) of complex traits in LD Hub ^11^, OpenGWAS database API, etc. We found that genes of the RelativeAge^(+)^ module overlapped significantly with top hits of GIANT Body fat distribution GWAS in humans (p<10^−3^, **TableS3, TableS7**).

**Fig 4.**
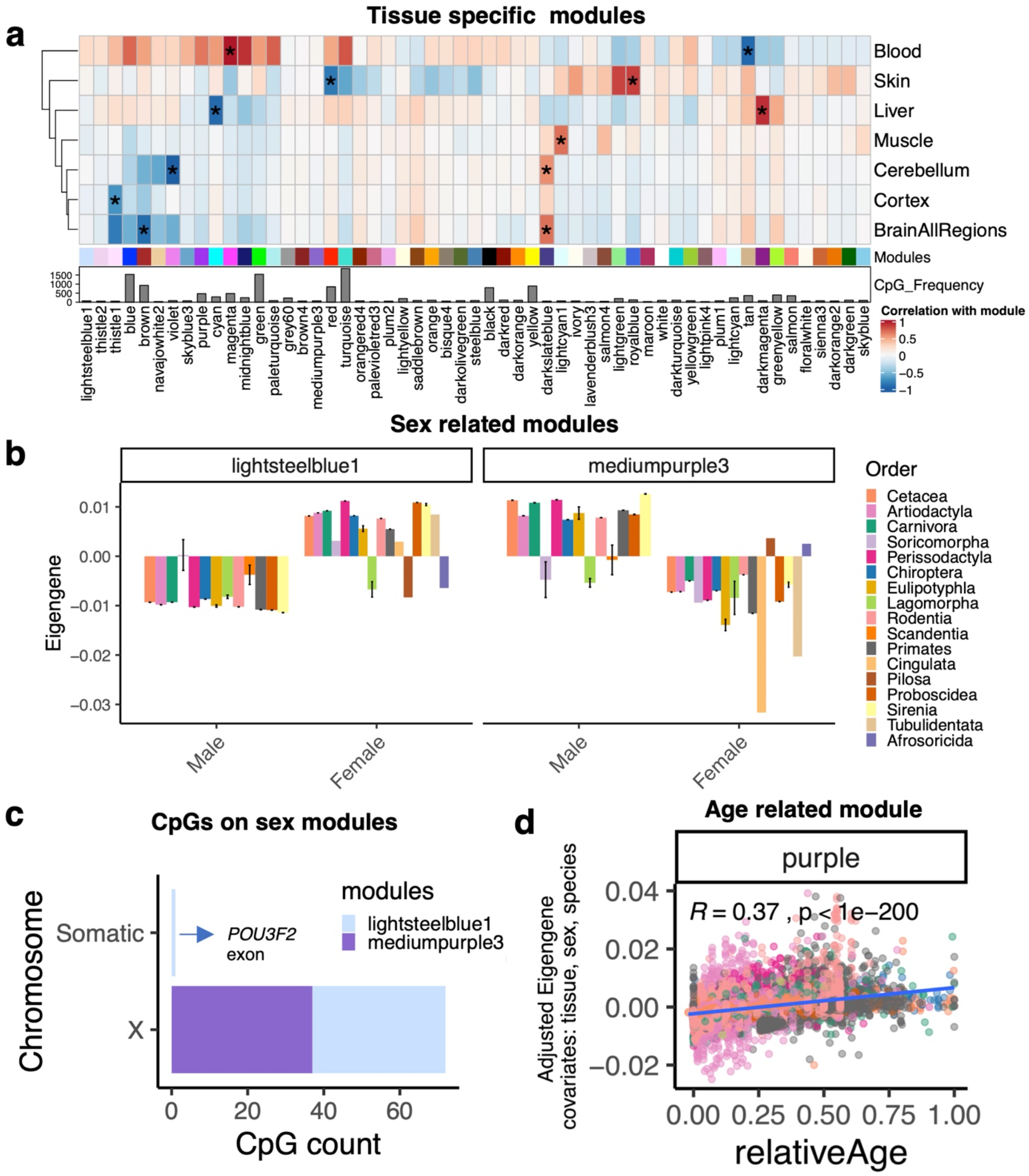
DNA methylation modules related to individual sample biological differences. **a**, the Heatmap of tissue specific modules. The tissues with no modules were excluded from the heatmap. * indicated the top two modules related to each tissue, cell type, or tissue region with minimum absolute correlation of 0.5. **b**, Sex-specific modules in different phylogenetic orders. **c**, Distribution of sex module CpGs on sex chromosomes. **d**, The top age-related module in all mammalian species (p<2.2e-16). The association with relative age (chronological age/maximum reported age) was examined after adjustment for tissue, sex and species differences. Eigengene: 1^st^ principal component of the module that positively correlates with DNA methylation levels.

### Life expectancy in pure dog breeds

As described above, five modules relate to maximum lifespan, adult weight, and age across most phylogenetic orders in our dataset, thereby allowing us to further investigate these associations in specific mammalian orders. Using DNA methylation data from n=574 blood samples from 51 different dog breeds (**Table S8**), we assessed whether these modules correlate with a measure of life-expectancy provided by the American Kennel Club ^12^. Interestingly, the LifespanWeight^(-)^Rodentia^(+)^ (greenyellow) module showed an inverse correlation with breed life-expectancy (r=-0.42, p<2×10^−16^) but a positive correlation with breed weight (r=0.37, p<2×10^−16^, **Fig. 5a**). The correlation pattern of this module is consistent with the well-known observation that larger dog breeds have shorter life expectancies ^13^. This finding is particularly interesting as this module also relates to the general mammalian trend where larger species have longer lifespans (Fig. 3a, 3b). The constituents and regulators of this module provide an entry point to understand the relationship between body size and lifespan. For example, as described above, LifespanWeight^(-)^Rodentia^(+)^ (greenyellow) module is partially related to retinoic acid receptor signaling and adipogenesis. Interestingly, a recent study showed that sphingomyelin levels are higher in large-short lived dogs ^14^. A substantially weaker correlation could be observed between the RelativeAge^(+)^ (purple) module and dog breed lifespan (r=-0.16, p=7×10^−5^).

**Fig 5.**
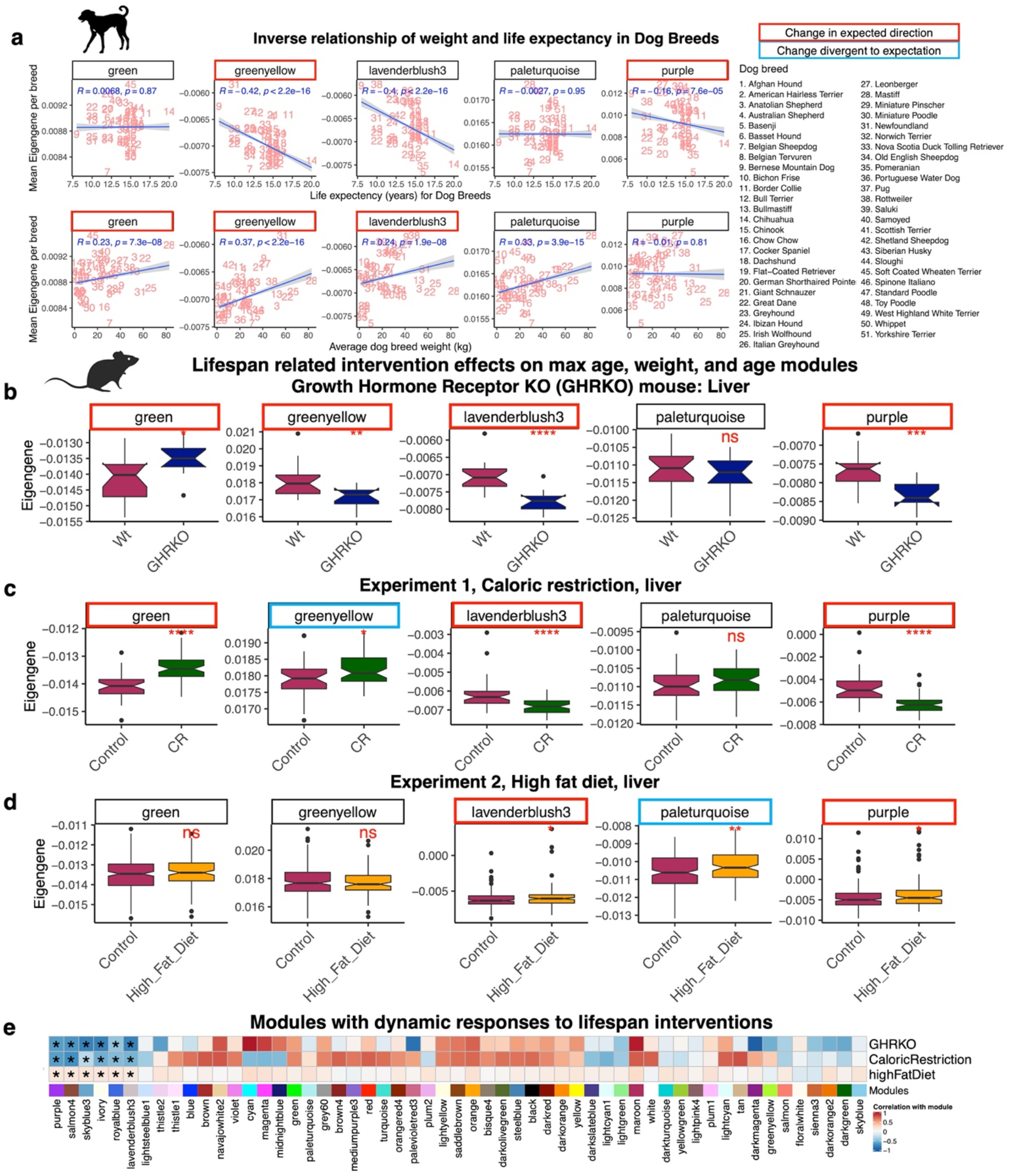
Defined DNA methylation modules are novel tools to study specific aging related and biological questions. **a**, DNA methylation modules corroborate the paradoxical inverse relation of life expectancy and average weight of dog breeds. Strikingly, the greenyellow module positively correlates with dog breeds’ weight, which is the opposite direction of the general pattern in mammalian species. **b**,**c**,**d**, DNA methylation modules are sensitive to lifespan-related intervention experiments and relate to the life expectancy of the animals. **b**, Changes in lifespan, weight, and age modules of liver parallels smaller size and longer life expectancy of growth hormone receptor mouse models (GHRKO). Sample size: GHRKO, 11 (5 female, 6 male); Wt, 18 (9 male, 9 female). The age range: 6-8 months. **c**, Caloric restriction (CR) DNA methylation module signature predicts longer lifespan in this treated group. This experiment only included male mice and the samples were collected at 18 months of age. Sample size: CR, 59; control, 36. **d**, High fat diet accelerates aging in the age module. Sample size: high fat diet, 133 (125 females, 8 males); control (*ad libido*), 212 (202 females, 10 males). Samples were collected throughout the lifespan from 3 to 32 months of age of both groups. **e**, Modules with dynamic responses to lifespan interventions. These modules are candidates for biomarkers of longevity. Only the modules with dynamic changes are labeled by ^*^ in the heatmap. ^*^ p<0.05; ^**^p<0.01; ^***^p<0.001; ^****^p<0.0001. Eigengene: 1^st^ principal component of the module that positively correlates with DNA methylation levels. Images of animals are from Phylopic (“http://phylopic.org“) or Wikimedia, which are under public domains or CC BY 3.0 license (“https://creativecommons.org/licenses/by/3.0/“).

### Interventional studies in mice

Our robust identification of epigenetic modules associated with traits such as age supports the notion that DNA methylation can serve as a dynamic molecular read-out of variations in individual-level phenotypes. Thus, we ascertained whether maximum lifespan, weight, and age modules are affected by interventions that are known to modulate the lifespan of mice. Growth hormone receptor knockout (GHRKO) mice, which have greatly increased lifespan ^15^, exhibited a change in specific modules: increase in LifespanWeight^(+)^Rodentia^(-)^ (green, p<0.05), decrease in LifespanWeight^(-)^Rodentia^(+)^ (greenyellow, p<0.01), decrease in Weight^(+)^ (lavenderblush3, p<0.0001), and decrease in RelativeAge^(+)^ (purple, p<0.001) eigengene values (**Fig. 5b**). The signs of these associations are consistent with the increased lifespan and smaller size of GHRKO mice ^15^. Caloric restriction (CR) and high fat diet are known to increase or reduce the lifespan of mice, respectively. Caloric restriction increased LifespanWeight^(+)^Rodentia^(-)^ (green, p<10^−5^), decreased Weight^(+)^ (lavenderblush3, p<0.0001), and decreased RelativeAge^(+)^ (purple, p<10^−5^) eigengenes in the expected direction (**Fig. 5c, d**). In contrast, CR moderately increased the LifespanWeight^(-)^Rodentia^(+)^ (greenyellow, p<0.05) eigengene in the opposite expected direction for longevity. High-fat diet was associated with an expected increase in the RelativeAge^(+)^ (purple, p<0.05) module but an unexpected increase in Lifespan^(+)^ (paleturquoise). Collectively, these studies demonstrate that some of the modules, especially those associated with lifespan, relative age, and weight across mammalian species, are indeed dynamic and can be reporters of anti-aging or pro-aging interventions within a species.

Additionally, we screened for any modules that showed a dynamic response to lifespan interventions. We identified a total of six modules with such a change. Future experiments would elucidate if these modules could serve as novel biomarkers of longevity and mortality risks in mammals.

## Discussion

This unbiased network analysis of the largest collection of DNA methylation data from 176 mammalian species facilitated the identification of methylation modules based on unsupervised clustering of highly-conserved CpGs. Thirty-one of the 55 modules could be interpreted by their associations with individual traits (chronological age, tissue, sex) or species traits (phylogenetic order, maximum lifespan, average adult species weight). The module-based analysis demonstrates that DNA methylation is a highly informative molecular read-out, not only at the level of individual tissues and within an organism, but also across species. This is evident from the high degree of congruence between the phyloepigenetic and phylogenetic trees. It indicates that species-level conservation and divergence of DNA methylation profiles closely parallels that of genetics through evolution. Moreover, several methylation modules show strong association with life history traits (e.g. maximum lifespan, average adult species weight) across 176 mammalian species, suggesting graded methylation changes on conserved DNA elements are robust features underlying the evolution of these quantitative traits in mammals. Overall, these results indicate that cytosine methylation data are highly informative for understanding the molecular basis of mammalian diversity. Although higher levels of DNA methylation are often associated with transcriptional silencing, a positive relationship between DNA methylation and expression levels has been observed especially in case of bivalent chromatin ^16^.

The disparities in aging rate and species lifespan are among the most intriguing biological mysteries that continue to engender debate over the relative importance of different aging theories, i.e. mutation accumulation ^17^, antagonistic pleiotropy ^18,19^, and disposable soma ^20^, which are predicated on a trade-off between reproductive fitness and longevity. Factors such as food availability, population density, and reproductive cost have been shown to impact this trade-off ^21^. This study supports the view that cytosine methylation relates to biological processes that underpin evolutionary differences in mammals ^22-25^.

While the identified modules lend themselves for studying many species traits, we were particularly interested in characterizing life history traits surrounding aging and development. In this context, we mention the RelativeAge^(+)^ (purple) module, which demonstrates conservation of DNA methylation aging biology across species and tissues. A decrease in the RelativeAge^(+)^ (purple) eigengene value, which is a collective decrease in the methylation level of CpGs in this module, related to an increase in life expectancy of dog breeds, GHRKO dwarf mice and caloric-restricted mice. In contrast, the high-fat diet increased the RelativeAge^(+)^ eigengene. Strikingly, the genes adjacent to the CpGs of this module are highly enriched for embryonic stem cell regulation, include targets of Polycomb repressor complex 2 (PRC2, EED) and H3K27me3 regions (**Extended Data Fig.7b**). These genomic features are strongly implicated in stem cell biology through key regulators such as SOX2 and NANOG transcriptional factors ^26,27^. This is corroborated by IPA analysis, which also highlighted NANOG and SOX2 as regulators of the RelativeAge^(+)^ (purple) module. Our findings extend similar findings from humans to other mammals ^28-30^, and demonstrates the intricate connection between stem cells, development and aging. Experimental perturbation of candidate regions in this module could elucidate if there is a causal link between module CpGs and lifespan of mammalian species. Future studies could evaluate whether module eigengenes associated with the RelativeAge^(+)^ (purple) and five other identified modules (**Fig. 5e**) lend themselves as indicators of biological age.

In summary, application of unsupervised machine learning through WGCNA-mediated clustering of CpG methylation has unveiled the stability of DNA methylation profiles at the species level while also identifying the responsiveness and dynamic nature of some methylation modules to anti-aging or pro-aging interventions in mice. The unsupervised approach provides a level of objectivity that gives a stamp of authenticity to the observation that DNA methylation is indeed a biologically meaningful molecular readout across all levels of mammalian life, from cells to species. The information-rich modules we have identified here form a roadmap to expand our approach towards cross-species DNA methylation analysis, and will undoubtedly open a new avenue to address many long-standing fundamental questions of biology, from evolution and lifespan to aging.

## Acknowledgment

This work was mainly supported by the Paul G. Allen Frontiers Group (SH). Nicolas Cermakian, Steve Brown and Stuart Peirson were involved in the mouse study of light pollution.

## Data Availability

The data will be made publicly available as part of the data release from the Mammalian Methylation Consortium. Genome annotations of these CpGs can be found on Github https://github.com/shorvath/MammalianMethylationConsortium

## Methods

### Data description

The data included 11,117 samples from 63 tissues of 176 mammalian species (167 eutherians, 9 marsupials). These samples were collected from different age ranges of most of the species. Sample collection and ethical approval for each mammalian species is described in separate individual papers ^8,31-47^. The species level characteristics such as maximum lifespan, average weight, and age at sexual maturity were chosen from anAge database ^48^. This data also includes DNA methylation from different dog breeds and mouse data from experimental lifespan intervention. Additional data sets included 57 horse transcriptome data generated from 29 different horse tissues ^46^. All DNA samples were analyzed by the novel custom-designed mammalian methylation array ^49^. This new array contains 38,608 probes that also includes 1,116 control probes. Following data collection, the SeSaMe normalization method was used to define beta values for each probe ^50^.

### Mappable CpGs for eutherians and mammals

The conserved probes were selected based on alignment of probes to 11 mammalian species from different phylogenetic orders ^49^. These species included Human (hg19), mouse (mm10), Vervet monkey (ChlSab1.1.100), Rhesus macaque (Mmul_10.100), Cattle (ARS-UCD1.2), Cat (Felis_catus_9.0.100), Dog (CanFam3.1), Elephant (loxAfr3.100), Bat (Rhinolophus_ferrumequinum.HLrhiFer5), Killer whale (GCF_000331955.2_Oorc_1.1), and Opossum (Monodelphis_domestica.ASM229v1.100). These species are selected based on a large sample size in our data, a relatively high genome quality, and also representation from different phylogenetic orders.

Two sets of probes were further selected from alignments in all these genomes. The first set was the probes that mapped to these ten eutherian species, and the second set were a subset that could also map to opossum as a marsupial representative. These two sets were additionally filtered using calibration data generated from the array’s performance on human, mouse, and rat synthetic DNA at different methylation levels (from 0-100% methylated) ^49^. Only the probes with a linear correlation of >=0.8 in all three species calibration data were kept as a mappable probe in mammalian species. The final number of remaining probes for the analysis were 14,705 in eutherians, and 7,956 that also mapped to marsupials.

### Unsupervised WGCNA

First, we formed two WGCNA networks based on the two sets of probes in our data. The first network was generated from 14,705 conserved CpGs in 10,939 samples of 167 eutherian species. The second network was a subset of 7,956 probes in 11,117 samples from 167 eutherians and 9 marsupial species. Traditionally, WGCNA is applied for transcriptome data and uses an unsupervised clustering method to assign the co-expressed genes into modules ^7^. In this study, we used the WGCNA method to define the modules of co-methylation CpGs in mammalian samples. First, the adjacency matrix (correlations between CpGs) was converted into a scale-free network using the soft threshold power (tuned value = 12) of the signed matrix. The result was converted into a topological overlap matrix (TOM), and 1-TOM distance measure (dissimilarity), which was used for Hierarchical clustering of the data. The trees were trimmed using a dynamic tree-cut algorithm to assign the modules containing at least 30 CpGs. Module eigengenes (MEs) were calculated as the maximum amount of the variance of the model that can be represented by a single variable for each module, based on the singular value decomposition method. The eigengenes in the eutherian network (Net 1) explained a range of 24-63% (average = 43%) of the variance in the methylation data in each module (**Table S3**). The hub CpGs of the modules were defined based on eigengene connectivity (kME) to each module. The association of module eigengenes were examined for different traits using multivariate linear regression models. The module colors in both networks were matched using the matchLabels() function in WGCNA package. Module preservation for each network was estimated using the “modulePreservation” R function in the WGCNA R package using primates as the reference for comparison.

### Consensus Networks

A total of seven consensus co-methylation networks were developed to effectively remove the confounding effects by conditioning on different species and tissue type combinations. The constructed consensus networks are as follows: cNet3, 57 species-tissue strata (network where tissue/species effects were removed); cNet4, 35 species but ignore tissue strata (only species effects were removed); cNet5, 15 tissue types but ignore species (only tissue effects were removed); cNet6, 27 species blood (species effects were removed in blood); cNet7, 7 species brain (species effects were removed in brain); cNet8, 10 species liver (species effects were removed in liver); and cNet9, 30 species skin data (species effects were removed in skin). Blood, skin, liver, and brain were tissues with data from >7 mammalian species.

Consensus WGCNA assumes that the DNA methylation network is conserved between multiple data strata. This network was generated following methods previously described ^7,51^. Briefly, the adjacency matrices (correlation) were constructed using DNA methylation beta values in each data set. The matrices were converted into scale free networks using a tuned soft threshold power for each dataset. Results were converted into TOM, merged, and then used to form a consensus tree network using a hierarchical clustering of dissimilarity matrix (1-TOM). Similarly, the colors were matched to network 1 colors.

### Hierarchical clustering

DNA methylation data from tissues with more than 50 species were used for hierarchical clustering and comparison with the evolutionary tree. The hierarchical clustering of tissue samples (as opposed to CpGs) was based on complete linkage coupled with a dissimilarity measure defined as 1-correlation. The distances in the hierarchical trees (i.e. the height values) were directly compared with the evolutionary distances (based on estimated time) in a publicly available evolutionary tree ^52^.

### Gene ontology enrichment

The genomic region level enrichment was performed using GREAT analysis ^53^ and the mappable probes as the background. The analysis used human hg19 annotations, a 50kb window for extending the gene regulatory domain, and default settings for the other options. For each module, the input included up to 500 hub CpGs. The biological processes were reduced to parent ontology terms using the “rrvgo” package. The larger ontology category was defined manually. The intra-module hub CpGs were also statistically tested for overlap with human GWAS results, in which gene p values were calculated by MAGENTA algorithm. Additionally, the hub genes were analyzed by ingenuity pathway analysis to identify the enriched canonical pathways and potential upstream regulators.

## Extended Data

**Extended Data Fig 1.**
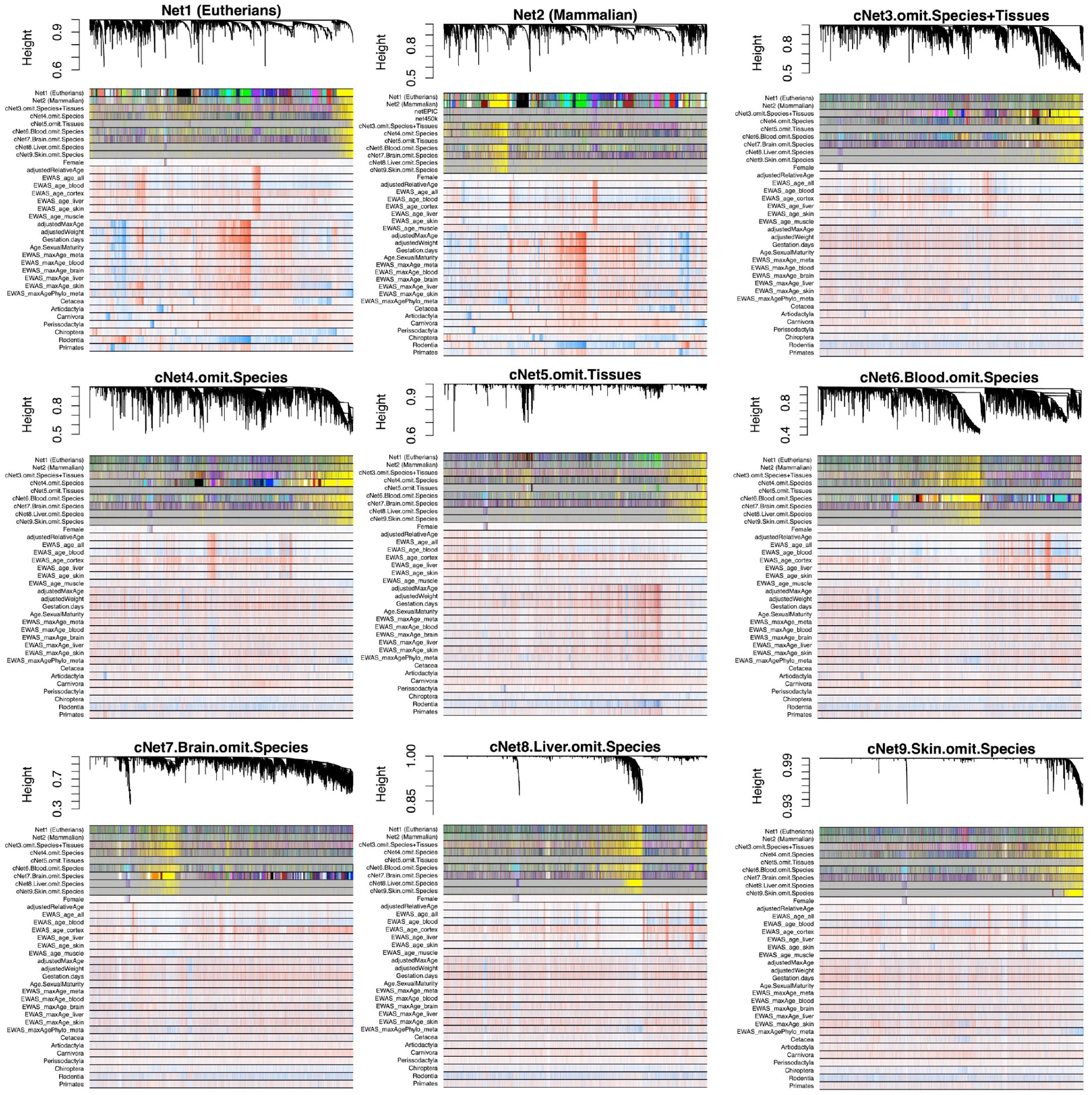
Constructed WGCNA networks from mammalian DNA methylation data. A total of nine WGCNA networks were constructed and compared for module association with species and individual sample characteristics. Network 1, an unsupervised network of 14,705 conserved CpGs in 167 eutherian species, showed the strongest module-trait associations. An additional unsupervised network was developed based on subsets of probes for future study applications: Network 2, 7,925 probes that map to both eutherians and marsupials. Additionally, seven consensus networks of 14,705 eutherian probes by different tissue and species combinations were formed to identify the most conserved modules for studying individual sample characteristics. All module colors were matched to network 1 for comparison.

**Extended Data Fig 2.**
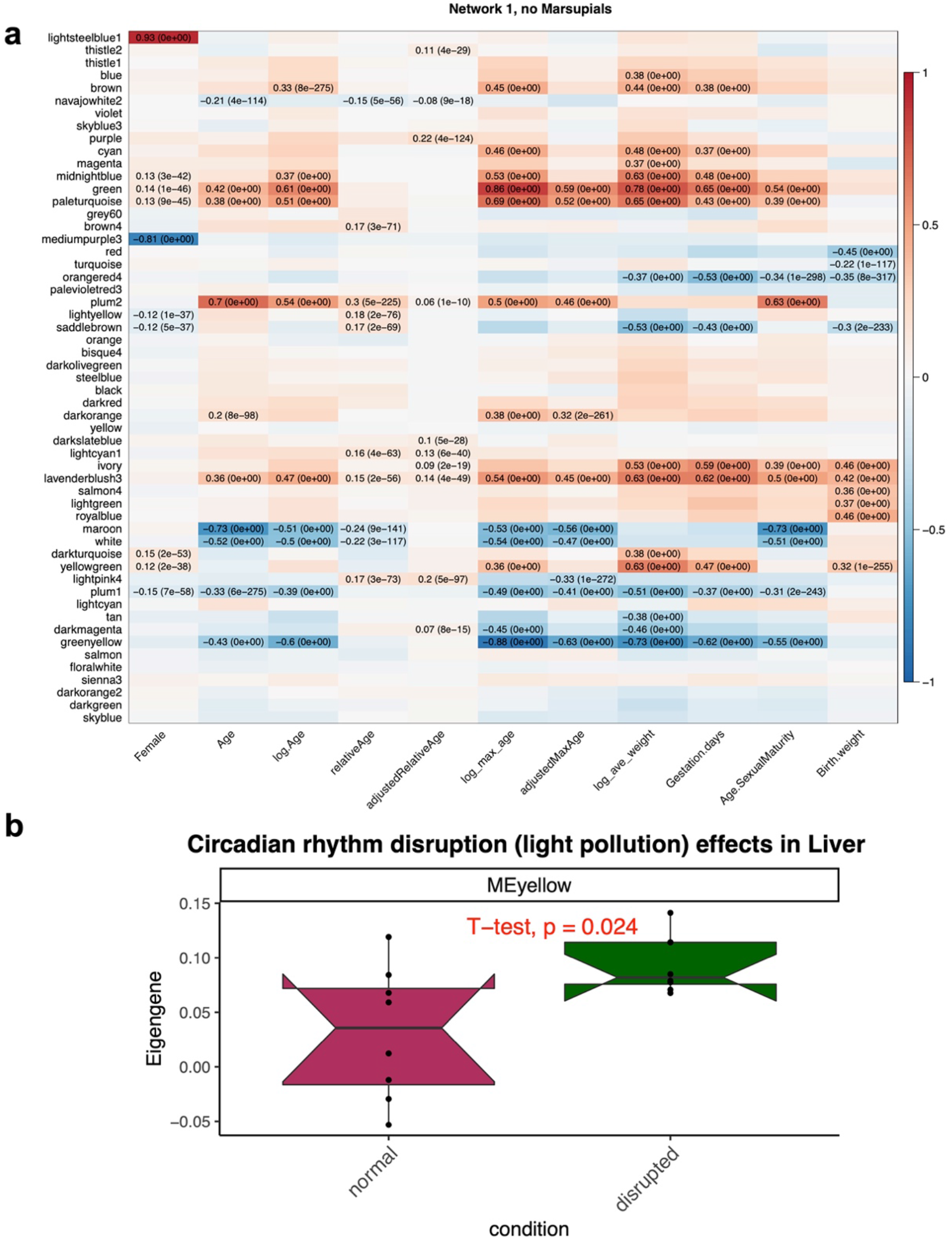
DNA methylation Module-trait association in mammals. **a**, Association of different individual and species-level traits with network 1 modules. Network 1 consists of 14,705 conserved CpGs in 167 eutherian species. The data include around 62 tissue types, from all age ranges of most of the species. **b**, Long term exposure to light pollution increases the yellow module eigengene. Control group, N = 8, standard light/dark cycles of 12 hours light (100 lux) followed by 12 hours dark. Circadian rhythm disrupted group, N = 8, the light cycle included12 hours light (100), followed by 12 hours dim light (20 lux). Cohorts were exposed to these conditions from three months of age, for a period of 12 months.

**Extended Data Fig 3.**
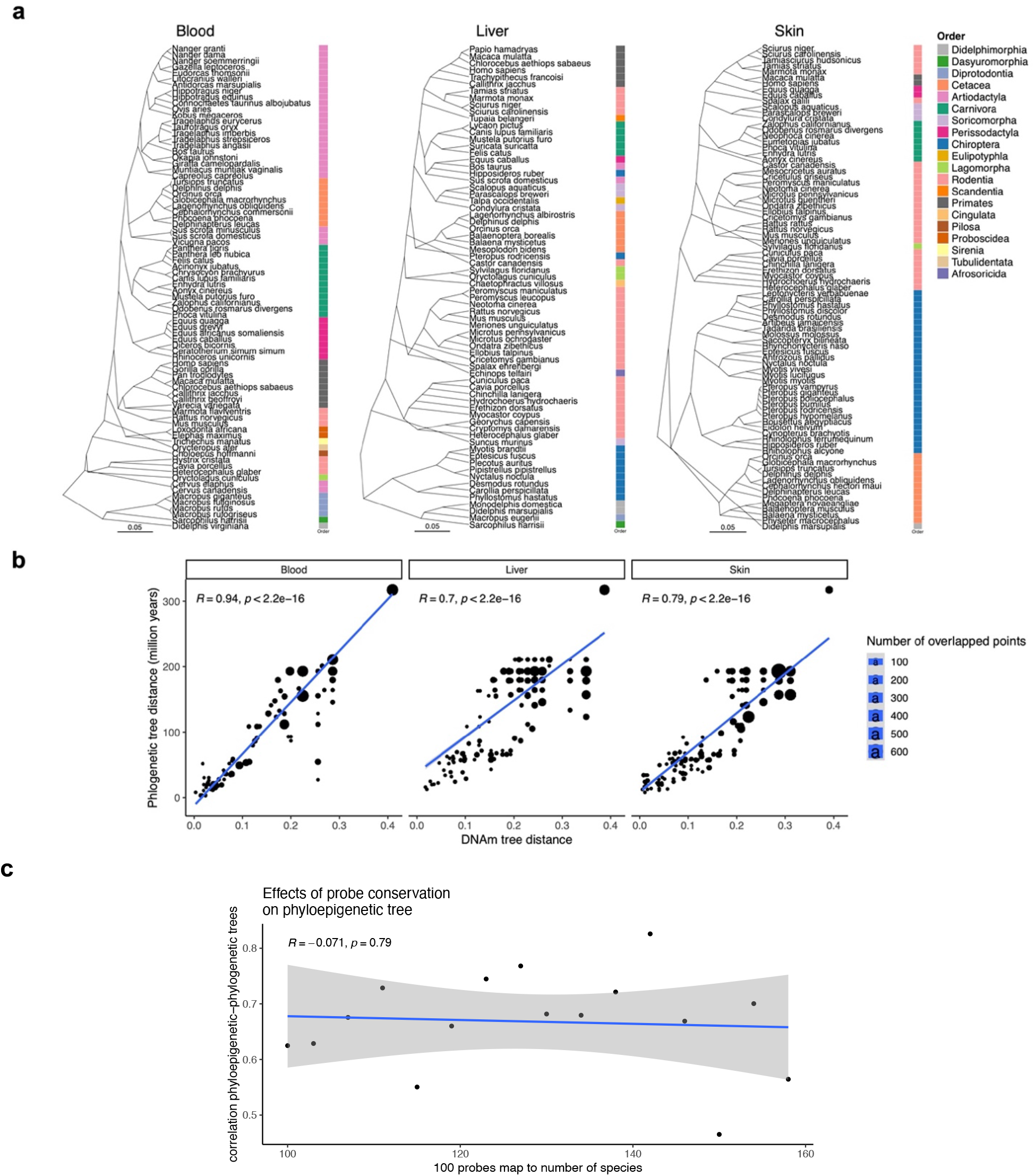
DNA methylation is a good indicator of mammalian evolution. **a**, Phyloepigenetic trees in blood, liver and skin of mammalian species. Distances: 1-cor. **b**, The distances of phyloepigenetic and evolutionary trees are highly associated. The size of the dots indicates the number of overlapping points in the plot. **c**, Sensitivity analysis of phyloepigenetic-phylogenetic trees relationship based on probe sequence conservation. 100 random probes were selected based on reduction in conservation and alignment to mammalian genomes. The phyloepigenetic distances (1-cor) were calculated from hierarchical clustering of blood DNA methylation data. The phylogenetic distances are based on the TimeTree database.

**Extended Data Fig 4.**
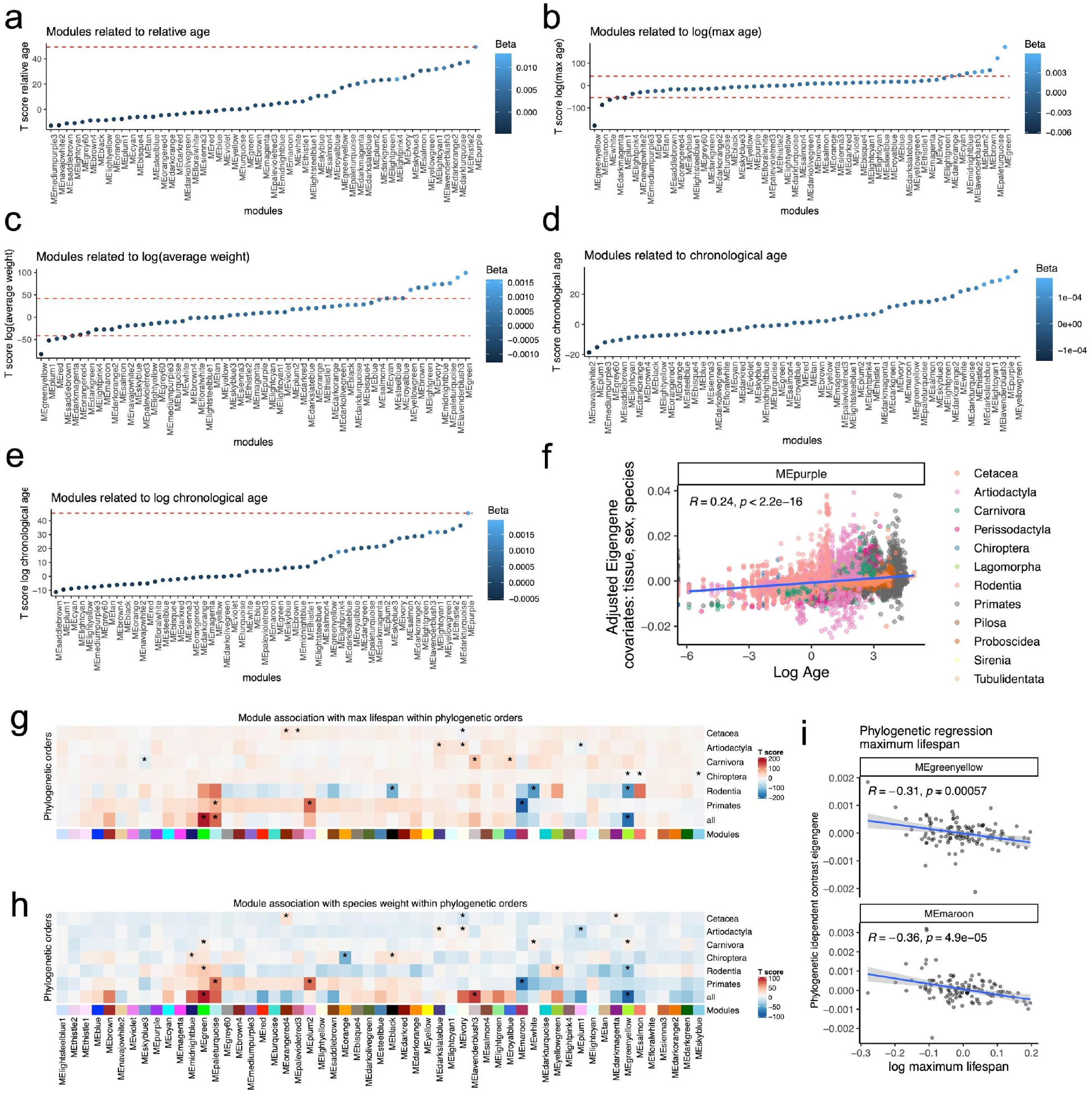
DNA methylation modules association with age, max age and weight in mammals. The association of modules eigengenes with relative age (**a**), log species maximum age (**b**), log species average adult weight (**c**), chronological age (**d**), and log chronological age (**e**). Red line: p =0. Covariates: tissue, sex and species differences for relative age, chronological age and log chronological age; relative age, tissue, and sex for log(max age) and log(average weight). **f**, Purple module also relates to log chronological age in mammals. Module association with log maximum age **(g)** and log species adult weight **(h)** within each phylogenetic order. ^*^ indicates the top 3 modules for max age, and weight modules for each order. **i**, Phylogenetic regression analysis of the modules with mammalian maximum lifespan. The evolutionary tree for phylogenetic regression was downloaded from the TimeTree database. Only the top two modules are presented in the scatter plot. Covariates: relative age, tissue, and sex.

**Extended Data Fig 5.**
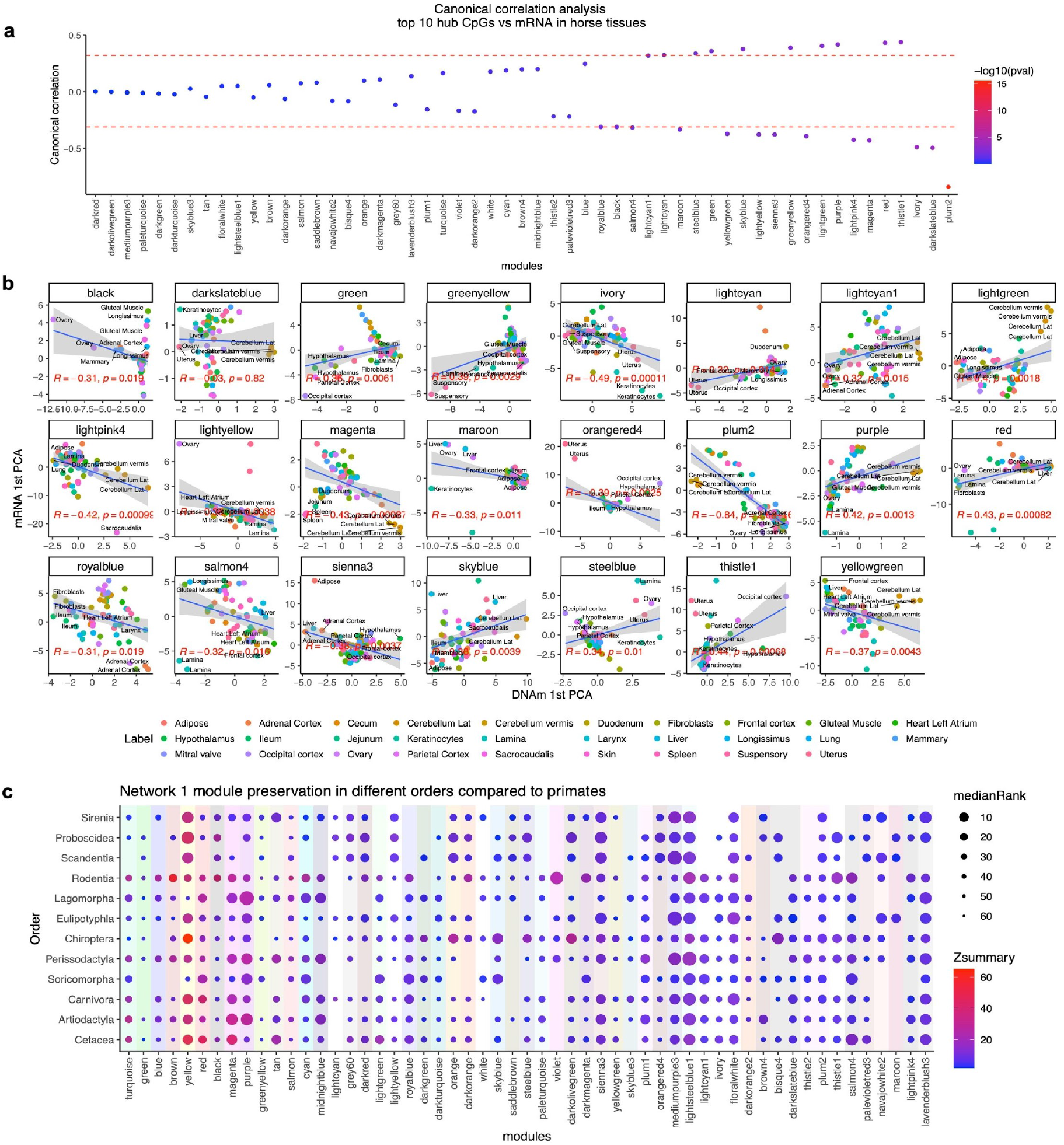
DNA methylation modules relate to mRNA changes and are preserved in different phylogenetic orders. **a**,**b**, DNA methylation modules and mRNA networks are canonically correlated in horse tissues. The mRNA and DNA methylation data originated from 57 samples from 29 different horse tissues. Canonical correlation is done for the 10 hub CpGs of each module and mRNA changes of their neighboring genes in the horse genome (EquCab3.0.100). The red lines indicate p<0.05. **c**, The DNA methylation modules are highly preserved in different phylogenetic orders. Module preservation is estimated with permutation of networks in each order and comparison with primates. The figures show the summary z score of preservation, summary p value of preservation, and median rank of preservation for each module.

**Extended Data Fig 6.**
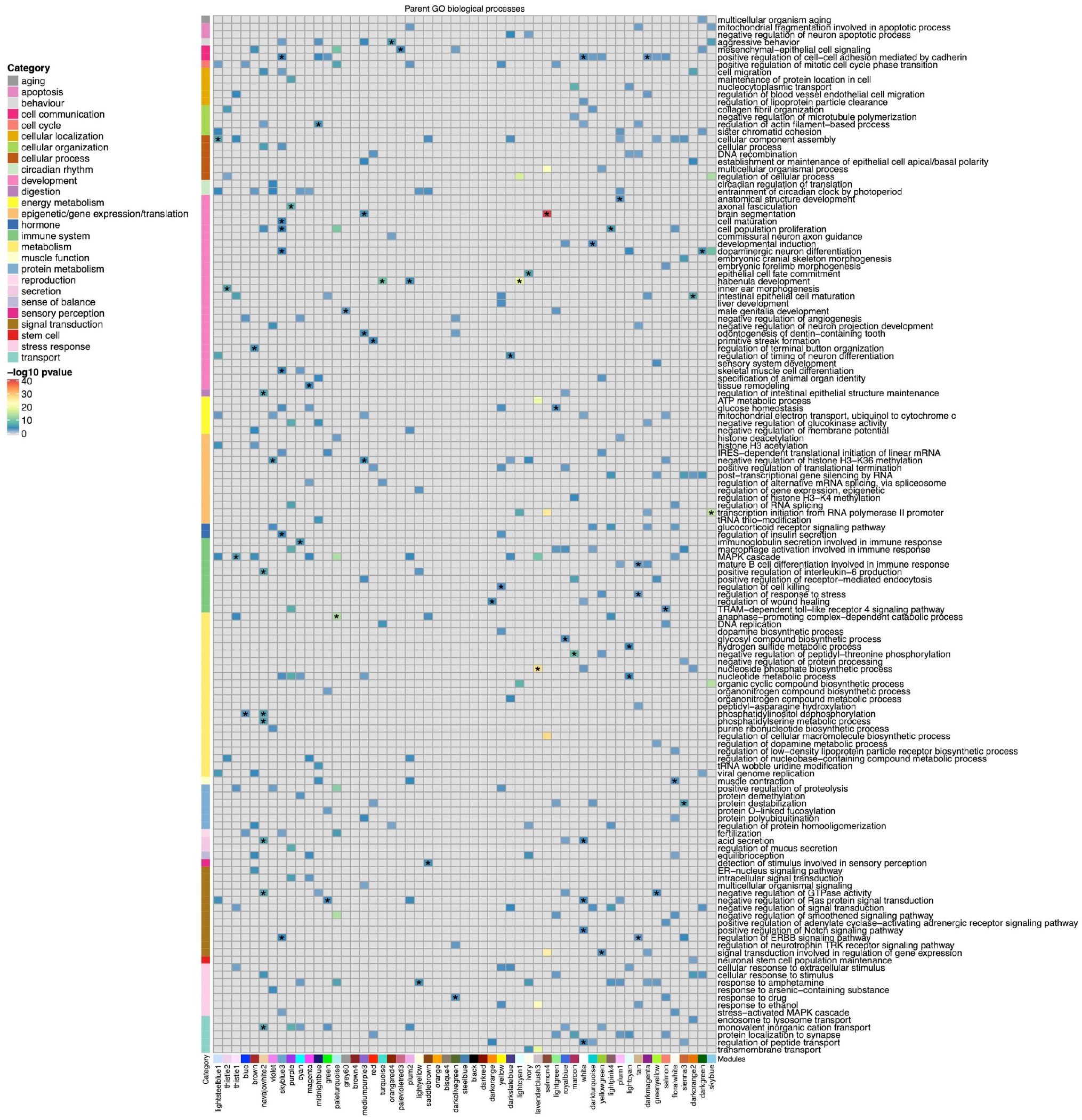
Gene ontology enrichment of the hub CpGs in each module. The gene level enrichment was done using GREAT analysis and human Hg19 background. The background probes were limited to 14,705 conserved probes in eutherians. The biological processes were reduced to parent ontology terms, and the top 10 parent terms with p<10^−3^ for each module were reported in the heatmap. The larger ontology category was defined manually and was used to arrange the ontology terms in the heatmap. ^*^ indicates the top ontology category for each module. The input includes the gene neighbors to up to 500 hub CpGs for each module.

**Extended Data Fig 7.**
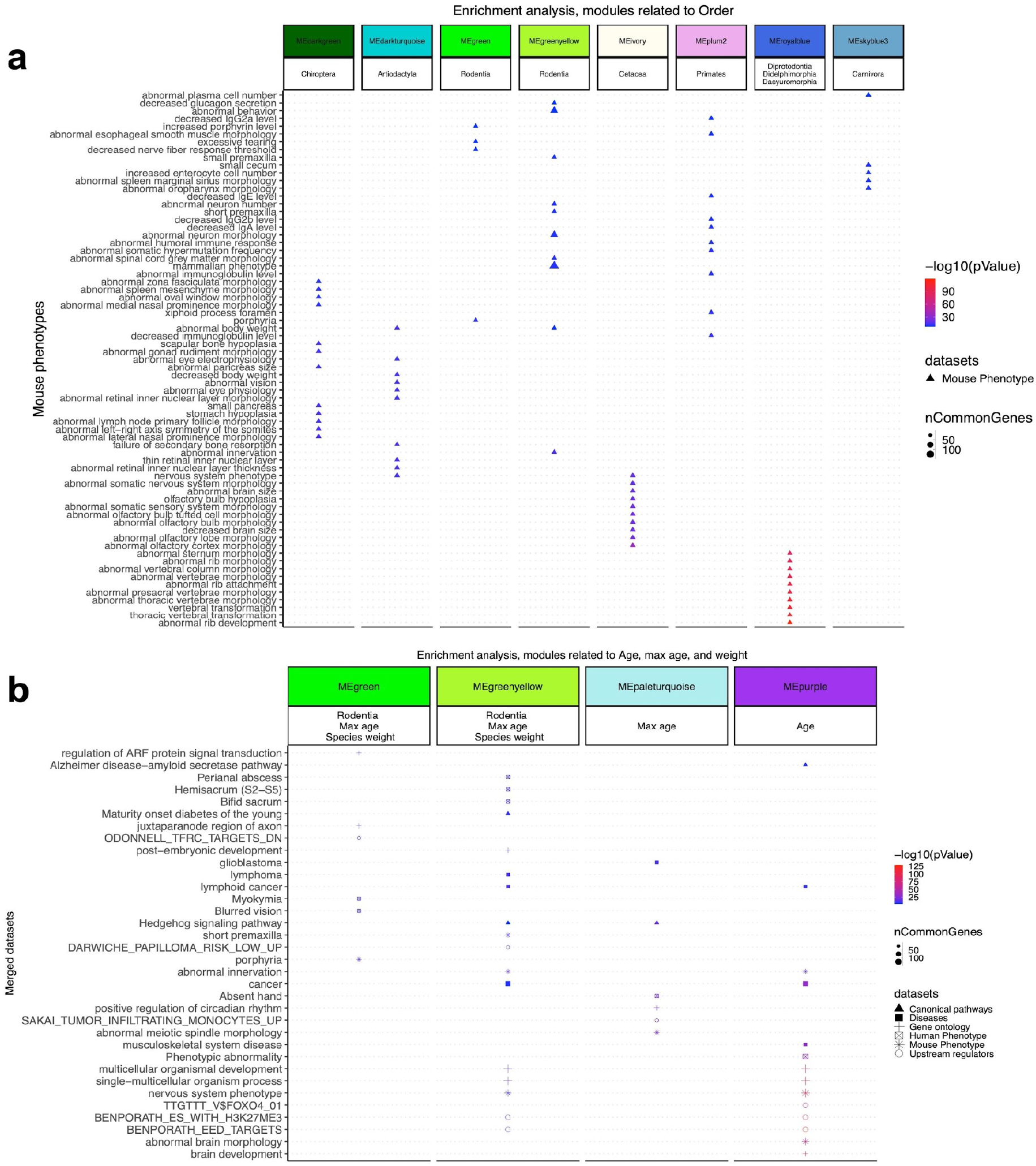
Enrichment analysis of modules associated with phylogenetic orders, age, max age, and average weight. The gene level enrichment was done using GREAT analysis and human Hg19 background. The background probes were limited to 14,705 conserved probes in eutherians. The input includes the gene neighbors to up to 500 hub CpGs for each module. **a**, Enrichment analysis of phylogenetic order modules. The plot shows the top unique enriched terms among the modules. **b**, Enrichment analysis of modules associated with max age and species weight. In addition to the unique enriched terms for each module, additional terms were mined based on specific keywords: ‘mortality’, ‘aging’, ‘survival’, ‘perinatal lethality’, and ‘weight’. The plot shows the top unique and top 20 overlapped enriched terms among the modules.

**Extended Data Fig 8.**
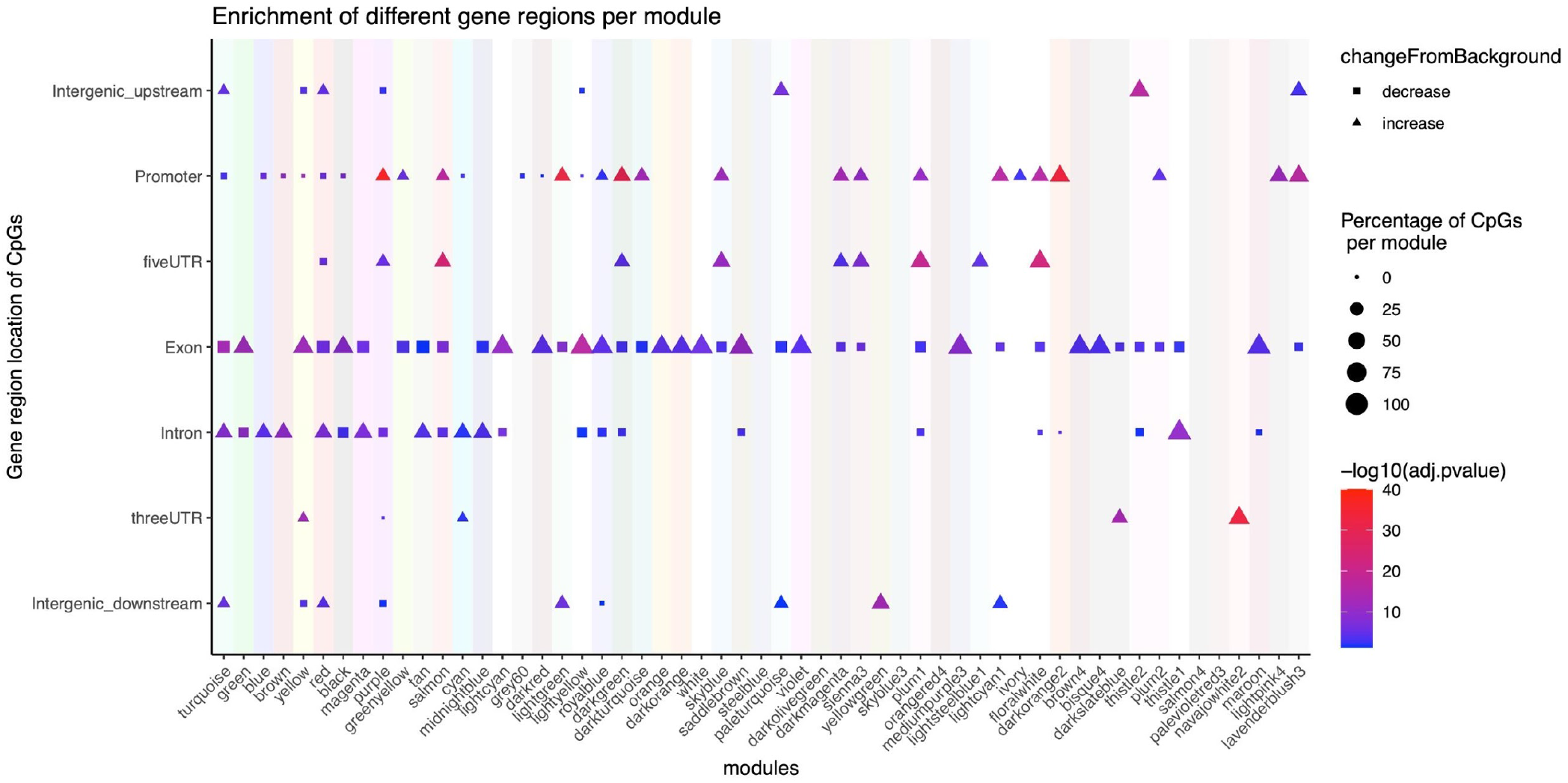
Enrichment of gene regions for CpGs in each module. The CpGs are annotated relative to the nearest transcriptional start site in the human Hg19 genome. The changes are estimated by proportion test (chi^2^) of each genomic region compared to the background. Only the regions with a significant difference than background are indicated in the dot-plot. The p-values are reported at 5% FDR for multiple test correction.

## References

1 John, R. M. & Surani, M. A. Genomic imprinting, mammalian evolution, and the mystery of egg-laying mammals. Cell 101, 585–588, doi:10.1016/s0092-8674(00)80870-3 (2000).

2 Welker, B. H. The history of our tribe: hominini. (2018).

3 Xiao, S. et al. Comparative epigenomic annotation of regulatory DNA. Cell 149, 1381–1392, doi:10.1016/j.cell.2012.04.029 (2012).

4 Villar, D. et al. Enhancer evolution across 20 mammalian species. Cell 160, 554–566, doi:10.1016/j.cell.2015.01.006 (2015).

5 Qu, J. et al. Evolutionary expansion of DNA hypomethylation in the mammalian germline genome. Genome research 28, 145–158, doi:10.1101/gr.225896.117 (2018).

6 Kimura, M. Evolutionary rate at the molecular level. Nature 217, 624–626, doi:10.1038/217624a0 (1968).

7 Langfelder, P. & Horvath, S. WGCNA: an R package for weighted correlation network analysis. BMC Bioinformatics 9, 559 (2008).

8 Horvath, S. et al. DNA methylation study of age and sex in baboons and four other primates. bioRxiv, 2020.2011.2029.402891, doi:10.1101/2020.11.29.402891 (2020).

9 Nasu, M. et al. Mammalian-specific sequences in pou3f2 contribute to maternal behavior. Genome Biol Evol 6, 1145–1156, doi:10.1093/gbe/evu072 (2014).

10 Lu, A. T. et al. Universal DNA methylation age across mammalian tissues. bioRxiv, 2021.2001.2018.426733, doi:10.1101/2021.01.18.426733 (2021).

11 Zheng, J. et al. LD Hub: a centralized database and web interface to perform LD score regression that maximizes the potential of summary level GWAS data for SNP heritability and genetic correlation analysis. Bioinformatics 33, 272–279, doi:10.1093/bioinformatics/btw613 (2016).

12 AmericanKennelClub. The Complete Dog Book: 20th Edition. (Ballantine Books, 2006).

13 Dammann, P., Sumbera, R., Massmann, C., Scherag, A. & Burda, H. Extended longevity of reproductives appears to be common in Fukomys mole-rats (Rodentia, Bathyergidae). PLoS One 6, e18757, doi:10.1371/journal.pone.0018757 (2011).

14 Hoffman, J. M., Kiklevich, J. V., Klavins, K., Valencak, T. G. & Austad, S. N. Alterations of Lipid Metabolism With Age and Weight in Companion Dogs. J Gerontol A Biol Sci Med Sci 76, 400–405, doi:10.1093/gerona/glaa186 (2021).

15 Pilcher, H. (Nature Publishing Group, 2003).

16 Bernhart, S. H. et al. Changes of bivalent chromatin coincide with increased expression of developmental genes in cancer. Sci Rep 6, 37393, doi:10.1038/srep37393 (2016).

17 Medawar, P. B. & University College, L. An Unsolved problem of biology; an inaugural lecture delivered at University College, London, 6 December, 1951. (H.K. Lewis and Co., 1952).

18 Williams, G. Pleiotropy, Natural Selection, and the Evolution of Senescence. Evolution 11, 398–411 (1957).

19 Austad, S. N. & Hoffman, J. M. Is antagonistic pleiotropy ubiquitous in aging biology? Evol Med Public Health 2018, 287–294, doi:10.1093/emph/eoy033 (2018).

20 Kirkwood, T. B. Evolution of ageing. Nature 270, 301–304 (1977).

21 Johnson, A. A., Shokhirev, M. N. & Shoshitaishvili, B. Revamping the evolutionary theories of aging. Ageing Res Rev 55, 100947, doi:10.1016/j.arr.2019.100947 (2019).

22 Lowe, R. et al. Ageing-associated DNA methylation dynamics are a molecular readout of lifespan variation among mammalian species. Genome biology 19, 22 (2018).

23 Maegawa, S. et al. Caloric restriction delays age-related methylation drift. Nature Communications 8, 539, doi:10.1038/s41467-017-00607-3 (2017).

24 Hernando-Herraez, I., Garcia-Perez, R., Sharp, A. J. & Marques-Bonet, T. DNA Methylation: Insights into Human Evolution. PLOS Genetics 11, e1005661, doi:10.1371/journal.pgen.1005661 (2015).

25 Verhoeven, K. J., vonHoldt, B. M. & Sork, V. L. Epigenetics in ecology and evolution: what we know and what we need to know. Mol Ecol 25, 1631–1638, doi:10.1111/mec.13617 (2016).

26 Lee, T. I. et al. Control of developmental regulators by Polycomb in human embryonic stem cells. Cell 125, doi:10.1016/j.cell.2006.02.043 (2006).

27 Boyer, L. A. et al. Polycomb complexes repress developmental regulators in murine embryonic stem cells. Nature 441, doi:10.1038/nature04733 (2006).

28 Teschendorff, A. E. et al. Age-dependent DNA methylation of genes that are suppressed in stem cells is a hallmark of cancer. Genome research 20, 440–446, doi:10.1101/gr.103606.109 (2010).

29 Rakyan, V. K. et al. Human aging-associated DNA hypermethylation occurs preferentially at bivalent chromatin domains. Genome research 20, 434–439, doi:10.1101/gr.103101.109 (2010).

30 Horvath, S. et al. Aging effects on DNA methylation modules in human brain and blood tissue. Genome Biol 13, R97, doi:10.1186/gb-2012-13-10-r97 (2012).

31 Wilkinson, G. S. et al. Genome Methylation Predicts Age and Longevity of Bats. bioRxiv, 2020.2009.2004.283655, doi:10.1101/2020.09.04.283655 (2020).

32 Sugrue, V. et al. Castration delays epigenetic aging and feminises DNA methylation at androgen-regulated loci. bioRxiv, 2020.2011.2016.385369, doi:10.1101/2020.11.16.385369 (2020).

33 Schachtschneider, K. M. et al. Epigenetic clock and DNA methylation analysis of porcine models of aging and obesity. bioRxiv, 2020.2009.2029.319509, doi:10.1101/2020.09.29.319509 (2020).

34 Sailer, L. L. et al. Pair bonding slows epigenetic aging and alters methylation in brains of prairie voles. bioRxiv, 2020.2009.2025.313775, doi:10.1101/2020.09.25.313775 (2020).

35 Raj, K. et al. Epigenetic clock and methylation studies in cats. bioRxiv, 2020.2009.2006.284877, doi:10.1101/2020.09.06.284877 (2020).

36 Prado, N. A. et al. Epigenetic clock and methylation studies in elephants. bioRxiv, 2020.2009.2022.308882, doi:10.1101/2020.09.22.308882 (2020).

37 Lemaître, J.-F. et al. Epigenetic clock and DNA methylation studies of roe deer in the wild. bioRxiv, 2020.2009.2021.306613, doi:10.1101/2020.09.21.306613 (2020).

38 Kordowitzki, P. et al. Epigenetic clock and methylation study of oocytes from a bovine model of reproductive aging. bioRxiv, 2020.2009.2010.290056, doi:10.1101/2020.09.10.290056 (2020).

39 Jasinska, A. J. et al. Epigenetic clock and methylation studies in vervet monkeys. bioRxiv, 2020.2009.2009.289801, doi:10.1101/2020.09.09.289801 (2020).

40 Horvath, S. et al. Reversing age: dual species measurement of epigenetic age with a single clock. bioRxiv, 2020.2005.2007.082917, doi:10.1101/2020.05.07.082917 (2020).

41 Horvath, S. et al. Epigenetic clock and methylation studies in the rhesus macaque. bioRxiv, 2020.2009.2021.307108, doi:10.1101/2020.09.21.307108 (2020).

42 Horvath, S. et al. DNA methylation age analysis of rapamycin in common marmosets. bioRxiv, 2020.2011.2021.392779, doi:10.1101/2020.11.21.392779 (2020).

43 Bors, E. K. et al. An epigenetic clock to estimate the age of living beluga whales. bioRxiv, 2020.2009.2028.317610, doi:10.1101/2020.09.28.317610 (2020).

44 Pinho, G. M. et al. Hibernation slows epigenetic aging in yellow-bellied marmots. bioRxiv, 2021.2003.2007.434299, doi:10.1101/2021.03.07.434299 (2021).

45 Horvath, S. et al. DNA methylation clocks show slower progression of aging in naked mole-rat queens. bioRxiv, 2021.2003.2015.435536, doi:10.1101/2021.03.15.435536 (2021).

46 Horvath, S. et al. DNA methylation aging and transcriptomic studies in horses. bioRxiv, 2021.2003.2011.435032, doi:10.1101/2021.03.11.435032 (2021).

47 Horvath, S. et al. Methylation studies in Peromyscus: aging, altitude adaptation, and monogamy. bioRxiv, 2021.2003.2016 (2021).

48 de Magalhaes, J. P., Costa, J. & Church, G. M. An analysis of the relationship between metabolism, developmental schedules, and longevity using phylogenetic independent contrasts. J Gerontol A Biol Sci Med Sci 62, 149–160 (2007).

49 Arneson, A. et al. A mammalian methylation array for profiling methylation levels at conserved sequences. bioRxiv, 2021.2001.2007.425637, doi:10.1101/2021.01.07.425637 (2021).

50 Zhou, W., Triche, T. J., Jr, Laird, P.W. & Shen, H. SeSAMe: reducing artifactual detection of DNA methylation by Infinium BeadChips in genomic deletions. Nucleic Acids Research 46, e123–e123, doi:10.1093/nar/gky691 (2018).

51 Langfelder, P. & Horvath, S. (UCLA, 2014).

52 Hedges, S. B., Marin, J., Suleski, M., Paymer, M. & Kumar, S. Tree of life reveals clock-like speciation and diversification. Mol Biol Evol 32, 835–845, doi:10.1093/molbev/msv037 (2015).

53 McLean, C. Y. et al. GREAT improves functional interpretation of cis-regulatory regions. Nat Biotechnol 28, doi:10.1038/nbt.1630 (2010).

54 Kumar, S., Stecher, G., Suleski, M. & Hedges, S. B. TimeTree: A Resource for Timelines, Timetrees, and Divergence Times. Molecular Biology and Evolution 34, 1812–1819, doi:10.1093/molbev/msx116 (2017).

